# Songbird subthalamic neurons signal song timing and error and project to dopaminergic midbrain

**DOI:** 10.1101/2021.08.30.458220

**Authors:** Anindita Das, Jesse H. Goldberg

**Author notes:** Correspondence, Address: W121 Mudd Hall, Cornell University, Ithaca, NY 14853. The authors declare no competing financial interests.

## Abstract

Skill learning requires motor output to be evaluated against internal performance benchmarks. In songbirds, ventral tegmental area (VTA) dopamine neurons (DA) signal performance errors important for learning, but it remains unclear which brain regions project VTA and how these inputs may implement the sensorimotor comparisons necessary for error computation. Here we find that the songbird subthalamic nucleus (STN) projects to VTA and that STN microstimulation can excite VTA neurons. We also discover that STN receives inputs from auditory cortical and ventral pallidal brain regions previously implicated in song evaluation. In the first neural recordings from songbird STN, we discover that the activity of most STN neurons is associated with body movements and not singing, but a small fraction of neurons exhibits precise song timing and performance error signals consistent with performance evaluation. Together our results implicate the STN-VTA projection as an evolutionarily conserved pathway important for motor learning and expand the territories of songbird brain associated with song learning.

**New & Noteworthy:** Songbird subthalamic (STN) neurons exhibit song-timing and performance error signals and are interconnected with auditory pallium, ventral pallidum and ventral tegmental area, three areas important for song learning.

## Introduction

Reinforcement learning (RL) relies on motor exploration and subsequent evaluation of performance outcome (Sutton and Barto 2018). Dopamine (DA) neurons in the ventral tegmental area (VTA) contribute to RL by sending reward prediction error signals to the basal ganglia (BG), where they regulate synaptic plasticity important for learning (Bromberg-Martin et al. 2010; Schultz et al. 1997). Phasic bursts in DA neurons reinforce recent actions while pauses promote extinction (Adamantidis et al. 2011; Tsai et al. 2009). While RL is canonically considered in the context of reward seeking, recent studies in songbirds suggest that common principles apply to motor performance learning, even when no external reward is at stake. Like motor skills in humans, song is a complex motor sequence learned through trial and error. Newborn zebra finches memorize a tutor song, or ‘template’, and as juveniles they practice, producing highly variable syllables akin to human babbling (Doupe and Kuhl 1999). Over weeks, variability decreases as the bird learns to produce a stereotyped syllable sequence resembling the tutor song (Tchernichovski et al. 2001). After tutoring, isolated juvenile zebra finches can learn to sing but deafened birds cannot, demonstrating that auditory-based self-evaluation is required for learning (Marler 1997). Social interactions may also play a role (Carouso-Peck and Goldstein 2019; Gadagkar et al. 2019).

Songbirds have a specialized neural circuit ‘the song system,’ that includes cortex-like nuclei necessary for normal song production and a DA-BG-thalamocortical loop necessary for song learning (Colquitt et al. 2021; Doupe et al. 2005; Murphy et al. 2017). Though the practicing bird does not receive explicit external rewards for hitting the right note, early theories of song learning proposed that practicing birds receive reinforcement signals for producing good song syllables (Doya and Sejnowski 1998). In support of this idea, recent studies show that song learning proceeds, at least in part (Hahnloser and Ganguli 2013), via an RL-like algorithm that is implemented in DA-BG circuits homologous to mammals (Chen et al. 2019; Fee and Goldberg 2011; Woolley 2019). First, songbirds can be trained to change the way they sing in response to syllable-targeted distorted auditory feedback (DAF) (Andalman and Fee 2009; Tumer and Brainard 2007). When a brief (50 ms) auditory distortion is targeted to specific variations of a targeted syllable, birds learn to change how they sing that syllable to avoid distortion (Tumer and Brainard 2007). Second, lesions to Area X, the BG nucleus of the song system, or its DA inputs impair both natural song learning as well as learning from DAF (Hoffmann et al. 2016; Sohrabji et al. 1990; Xiao et al. 2018). Third, Area X-projecting DA neurons exhibited phasic suppressions following DAF distortions, and activations at the precise moment of the target syllable when a predicted distortion did not occur (Gadagkar et al. 2016), consistent with an RPE-like signal (Schultz 1998). Finally, optogenetic activation of DA axons in Area X reinforces syllable renditions (Hisey et al. 2018; Xiao et al. 2018). Together, these studies show that birds learn to sing using circuits that are also known to play roles in reward seeking during foraging (Chen and Goldberg2020).

The discovery of RPE-like signals in DA neurons during singing raised the question of how inputs to VTA convey information about song quality. Anatomical studies showed that VTAx neurons receive inputs from at least two forebrain inputs: an auditory cortical region (AIV, ventral anterior intermediate arcopallium) and a ventral pallidal (VP) region (Gale et al. 2008; Mandelblat-Cerf et al. 2014; Mello et al. 1998). Follow-up electrophysiological investigations clarified roles of AIV and VP in song evaluation. First, VTA projecting AIV neurons (AIVvta) exhibit errorinduced activations, AIV stimulation drives suppressions in VTAx firing (Chen et al. 2019; Mandelblat-Cerf et al. 2014), and syllable-targeted optogenetic activation of AIV axons in VTA extinguish syllable variations (Kearney et al. 2019; Xiao et al. 2018). These findings all suggest that AIV ‘error’ signals drive pauses in VTA following worse-than-predicted syllable outcomes, which in turn extinguish those variations. Meanwhile, VP neurons exhibit diverse error and predicted error signals during singing, VP stimulation can drive activation of VTAx neurons, and photoactivation of VP axons in VTA reinforces syllable variations (Chen et al. 2019; Kearney et al. 2019; Xiao et al. 2018). These findings support the idea that VP sends predicted error signals to VTA that can drive bursts when predicted distortions do not occur.

Yet one shortcoming of this model is that the VPvta neurons thought to drive DA bursts are mostly GABAergic, such that DA bursting would have to result from a post-inhibitory-rebound (PIR) mechanism. While PIR can drive DA activations in vitro (Paul and Johnson 2003), mice with NMDA receptors knocked out of DA neurons exhibit impaired learning and impaired DA bursting, revealing the importance of glutamatergic inputs for DA burst generation (Zweifel et al. 2009). The absence of an excitatory input to DA neurons in songbirds that are poised to drive phasic bursts during singing led us to hypothesize that excitatory inputs to songbird VTA from the subthalamic nucleus (STN) both exist and may play a role in singing. Two recent studies inspired this idea. First, the STN was recently shown to be interconnected with the same part of the songbird VP that sends error signals to VTA (Jiao et al. 2000; Person et al. 2008). Second, a landmark study in mice that used rabies virus-assisted whole brain mapping to DA neurons revealed, surprisingly, that the STN is the dominant source of monosynaptic excitation to midbrain DA neurons (Tian et al. 2016). This projection gives a functionality to reward and reward prediction error signals previously observed in STN (Baunez et al. 2005; Darbaky et al. 2005; Lardeux et al. 2009) – signals that did not support more mainstream models proposing that the STN primarily functions as part of the hyperdirect pathway and serves to provide a ‘brake’ on ongoing movements due to its strong connection to GABAergic outputs of the BG (Fife et al. 2017; Nambu et al. 2002; Schmidt et al. 2013; Wessel et al. 2016).

We thus set out to test if the songbird STN, which is located outside the classic song system and which has not previously been studied in the context of singing, may exhibit singing-related functions. We find, first, that the STN receives inputs from the VTA-projecting parts of both AIV and VP, which could enable sensorimotor comparisons between ‘actual’ and ‘predicted’ auditory feedback during singing. Second, the STN projects to and modulates the discharge of VTA neurons. Finally, we recorded STN neurons in singing birds for the first time and discovered a small, scattered fraction that exhibits precise syllable timing and performance error signals. Together these findings show that, as in mammals, the STN receives cortical inputs, is reciprocally connected with VP, and projects to VTA. Given the evolutionarily conserved subcortical circuitry in songbirds and mammals (Jiao et al. 2000; Person et al. 2008), our findings suggest that the STN may play a general role in performance error computations important for learning.

## Materials and Methods

### Subjects

Subjects were 52 adult male zebra finches (at least 90 days post hatch, dph). Animal care and experiments were carried out in accordance with NIH guidelines and were approved by the Cornell Institutional Animal Care and Use Committee.

### Surgery and histology

For all surgeries, birds were anaesthetized under isofluorane. All anatomical co-ordinates are expressed as antero-posterior (A) and medio-lateral (L) direction with respect to the lambda and dorso-ventral (D/V) direction with respect to the pial surface. To test if STN projected to VP, 100nl of green fluorescent protein (GFP)-expressing self-complementary adeno-associated virus (scAAV9-CBh-GFP, UNC vector core) was injected into VP (4.9A, 1.3L, 3.8V at 20 degrees head angle) bilaterally. To identify areas projecting to STN, 4 birds were injected with 100nl scAAV9-CBh-GFP bilaterally in STN at co-ordinates corresponding to the cluster of VP-projecting neurons in the anterior diencephalon (4.6A, 0.6L, 4.6/4.9V at 40 degrees head angle). In 2 of these birds, 50nl of fluorescently labeled cholera toxin subunit B (CTB, Molecular Probes) was also injected bilaterally in Area X (5.6A, 1.5L, 2.65V at 20 degrees head angle) to test for co-labeling in areas that projected to both STN and Area X. In 2 other birds, 50nl CTB was injected bilaterally in VTA (2.4A, 0.6L, 6.3V at 55 degrees head angle) to examine co-labeling in areas that projected to both STN and VTA. In another set of 2 birds, 70nl CTB was injected in STN bilaterally (4.2/4.6A, 0.6L, 4.6/5.0V at 40 degrees head angle) to confirm the results from retrograde viral injections. In a set of 2 birds, ~40nl anterograde virus HSV-mCherry (MGH, Viral Core) was injected in STN along with CTB in Area X to test if STN axons localized with X-projecting VTA neurons. Birds were perfused with 4% paraformaldehyde solution and 100 μm sagittal slices were obtained for imaging. All imaging was acquired using Leica DM4000 B microscope except for imaging STN axons in VTA (Fig. 1D) that was obtained using Zeiss LSM 710 Confocal microscope.

**Fig. 1.**
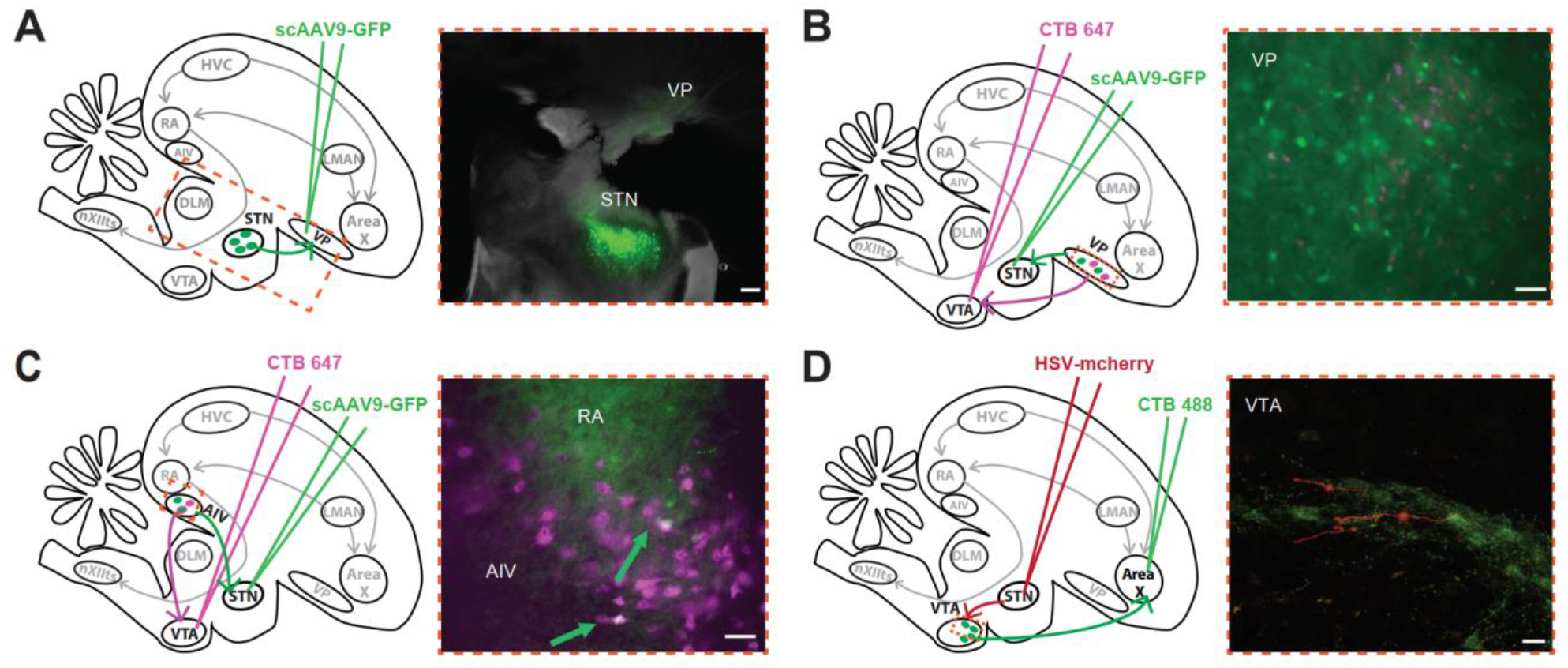
STN is reciprocally connected to VP, receives projections from AIV and sends projections to VTA neurons. A. Injection of retrograde viral tracer scAAV9-GFP in VP resulted in labelled cells in STN, confirming its anatomical location in zebra finches (Scale bar: 250 μm). B. Injection of scAAV9-GFP in STN and retrograde tracer, choleratoxin subunit B 647 in VTA were performed to test for collateral projections from VP to STN and VTA. STN-projecting cells were observed intermingled with VTA-projecting cells in VP, without colabelling. (Scale bar: 50 μm) C. Injection of scAAV9-GFP in STN and retrograde tracer, choleratoxin subunit B 647 in VTA were performed to test for collateral projections from AIV to STN and VTA. Collabelled cells were observed in AIV (green arrows) suggesting the presence of a population of AIV neurons sending projections to both STN and VTA. (Scale bar: 50 μm) D. Anterograde viral tracer HSV-mCherry was injected in STN along with retrograde CTB-488 in AreaX to assess if STN projected to X-projecting VTA neurons (VTAx). Image was obtained under confocal microscope, which showed labelled axons from STN adjacent to VTAx neurons (Scale bar: 20 μm).

### Functional mapping and neural analysis

For functional mapping experiments, birds (*n*=8) were implanted with bipolar stimulating electrodes in Area X and STN. All recordings were performed in anaesthetized birds. VTA was identified with reference to the DLM-Ovoidalis boundary (Gadagkar et al. 2016) and VTA neurons were recorded using a carbon fiber electrode (1 MOhm, Kation Scientific). Area X-projecting VTA neurons (VTAx) were confirmed by antidromic response and collision testing (200 μs pulses, 100-300 μA) using the Area X stimulating electrode. All VTA neurons recorded were tested for response to STN stimulation. Small electrolytic lesions were made by passing ±30 μA of current for 60 sec through the electrodes at the end of the experiment to confirm the locations of stimulation and recording. The recorded VTA neurons (*n*=13) were analyzed using custom software in MATLAB. Spike sorting was performed and spike rasters aligned to stimulation times were computed to assess the response of VTA neurons to STN stimulation. Firing rate histograms were computed using 2 ms bins to account for the short latencies to response of the VTA neurons to STN stimulation and smoothed with a 3-bin moving average. Each bin was tested for significant firing rate changes using a z-test (*p* < 0.05). Latency to response was defined as the first bin for which the next 2 consecutive bins were significantly different from previous activity (z-test, *p* < 0.05). Duration of response was computed from the total number of significant bins.

### Awake-behaving electrophysiology

For awake-behaving electrophysiology, birds (*n*= 36) were implanted with 16 channel movable electrode bundles (Innovative Neurophysiology) over STN (4.1-4.7A, 0.6L, 4.2V). Birds with 16 channel movable implants were placed in sound isolation chambers, maintaining a 12 hr light-dark cycle. They were allowed to recover for a day post-op and then subjected to distorted auditory feedback (DAF) protocol to habituate them before starting neural recording. Real-time analysis of singing was carried out using custom-written acquisition program in LabView to deliver ~25 ms DAF on top of a specific syllable during the song 50% of the times (Gadagkar et al. 2016). DAF was broadband noise filtered at 1.5-8 kHz to match the spectral range of the finch song and delivered through dual speakers placed inside the recording chamber. At the end of the experiment, small electrolytic lesions were made to ascertain the location of the recording site.

### Neural recording and analysis

Neural recordings were obtained using 16 channel INTAN headstages with the accompanying recording controller and INTAN acquisition software at a sampling rate of 20kHz. The song and a copy of the DAF signal was recorded contiguously with the neural data through analog inputs on the INTAN recording controller to facilitate time alignment of all data. The neural data was analyzed further using custom software in MATLAB. Spike sorting was performed and firing rate histograms were computed using 10 ms bins and smoothed with a 3-bin moving average, except for analysis of the song-locked neurons that were computed with 4 ms bins (Fig. 3A-C). To further quantify the firing of neurons locked to song timing, inter-motif correlation coefficient (IMCC) was calculated as described previously (Chen et al. 2019; Kao et al. 2008; Ölveczky et al. 2005). Movement analysis was performed on data from a subset of birds (*n*=4) that were acquired using 16 channel INTAN headstages mounted with a 3-axis accelerometer. The net acceleration was computed as follows:

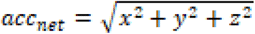

where, *x*, *y* and *z* are the acceleration values from the 3-axis accelerometer signal. This net acceleration was used to compute thresholds for movement onsets and offsets using k-means clustering (Chen et al. 2019; Gadagkar et al. 2016). Movement onsets and offsets were detected based on threshold crossings of the smoothed (moving average with 25 ms windows) rectified net acceleration signal. Firing rate histograms were then computed using 10 ms bins and smoothed with a 3-bin moving average. To test for significant movement-related firing rate changes, firing rate histograms were computed for ±300 ms of spiking activity around the movement onsets or offsets with 10 ms binsize and steps of 5 ms. Each bin was tested for significance using a z-test (*p* < 0.05). Latency of response was computed based on the first bin (w.r.t movement onset/offset) for which four consecutive bins were significant. We further classified movement events as during singing and outside singing to assess if the neurons encoded movement differentially during the two behavioral states.

## Results

### STN receives inputs from Auditory pallium and Ventral Pallidum in zebra finches

In mammals, the STN receives topographically organized inputs from widespread cortical, BG, and limbic areas, including VP (Joel and Weiner 1997). Past work in pigeons and songbirds identified a small area in the zebra finch diencephalon that is reciprocally connected to the VP, an anatomical signature for STN (Groenewegen et al. 1993; Haber et al. 1993; Jiao et al. 2000; Person et al. 2008). Here, we confirmed this anatomical connection. Injection of retrograde virus into VP resulted in labeling of a small area in the diencephalon, ~ 0.8 mm anterior to Ovoidalis and 1-0.8 mm ventro-medial to VP (2/2 hemispheres) (Fig. 1A). Retrograde injections in STN also resulted in back-labeled neurons into the part of VP that we and others recently showed projects to VTA (8/8 hemispheres) (Fig. 1B) (Chen et al. 2019; Gale et al. 2008; Kearney et al. 2019). To test if VP-STN neurons are collaterals of VP-VTA neurons, we injected retrograde tracers into both STN and VTA into single hemispheres (*n*=4 hemispheres) and failed to observe co-labeling, suggesting that independent populations of VP neurons project to STN and VTA (4/4 hemispheres). Interestingly, following STN retrograde viral injections, we also discovered labeled cells in the part of AIV that projects to VTA (2/4 hemispheres). To test if AIV neurons that project to VTA also project to STN, we examined AIV neuron labeling in hemispheres doubly injected with retrograde tracer in STN and VTA. Co-labeled cells were observed in AIV, suggesting that a population of AIV neurons that projects to VTA also projects to STN (Fig. 1C). We additionally confirmed these results by injecting CTB into STN that resulted in a larger population of cells being labeled in the AIV (not shown here, 2/2 hemispheres). Past work showed that both AIV and VP project to VTA and exhibit error signals during singing (Chen et al. 2019; Mandelblat-Cerf et al. 2014). The discovery that both these areas also project to STN raises the possibility that the STN could also play a role in song learning.

### Songbird STN projects to VTA and activates VTA neurons

We next wondered if songbird STN projects to the part of VTA that projects to Area X. We injected anterograde virus HSV-mCherry in STN and CTB in Area X to label STN axons and Area X-projecting neurons, respectively. We observed labeled axons from STN near VTAx neurons (Fig. 1D). Imaging with confocal microscopy further allowed us to visualize axon boutons from STN in close proximity to VTAx neurons and dendrites (Fig. 1D). While confirmation of actual synaptic contacts would require electron microscopy, these anatomical findings are consistent with the known STN-DA connectivity in mammals (Joel and Weiner 1997; Kooy and Hattori 1980; Ogawa et al. 2014; Watabe-Uchida et al. 2012), consistent with general conservation of BG anatomy across vertebrates (Grillner et al. 2013; Reiner et al. 1998).

To test for functional connectivity between STN and VTA, we performed mapping experiments in anaesthetized birds (*n*=8 birds) (Fig. 2A). A recording electrode was used to record VTA neurons while stimulation electrodes in Area X and STN were used for antidromic identification and orthodromic responses in VTA neurons, respectively. Stimulation of Area X was used to antidromically identify Area X-projecting VTA neurons, confirmed by collision testing (*n*=7) (Fig. 2B). VTA neurons that did not exhibit antidromic response to Area X stimulation were categorized as VTAother neurons (*n*=6). All VTA neurons were tested for response to STN stimulation. STN stimulation mostly caused activations (Fig. 1C), including in VTAx neurons (Fig.1D). Neurons that exhibited activation to a single pulse of STN stimulation were further tested with bursts of STN stimulation at subthreshold intensities to distinguish between antidromic and orthodromic response. By this criteria and the absence of any observed antidromic collisions, we concluded that the response of the VTAx neurons to STN stimulation, though highly short latency and precise, were orthodromic, though we cannot rule out antidromic responses as neuronal firing was too sparse for collision testing. Most VTAx neurons exhibited a short latency (2.5 ± 0.61 ms, mean ± s.d,) activation to STN stimulation (*n*=5/7); two neurons exhibited suppressions (example in Fig. 1E). All VTAother neurons exhibited a short latency (mean ± sd, 3.25 ± 1.13 ms) activation to STN stimulation (example in Fig. 1F). Together, these results suggest that STN projects to VTA and can modulate the firing of VTA neurons, including VTAx neurons that have been shown to exhibit performance error signals in singing birds. Yet we cannot rule out the possibility that the STN stimulation also activated fibers of passage coursing through the diencephalon en-route to VTA, including projections from AIV and VP.

**Fig. 2.**
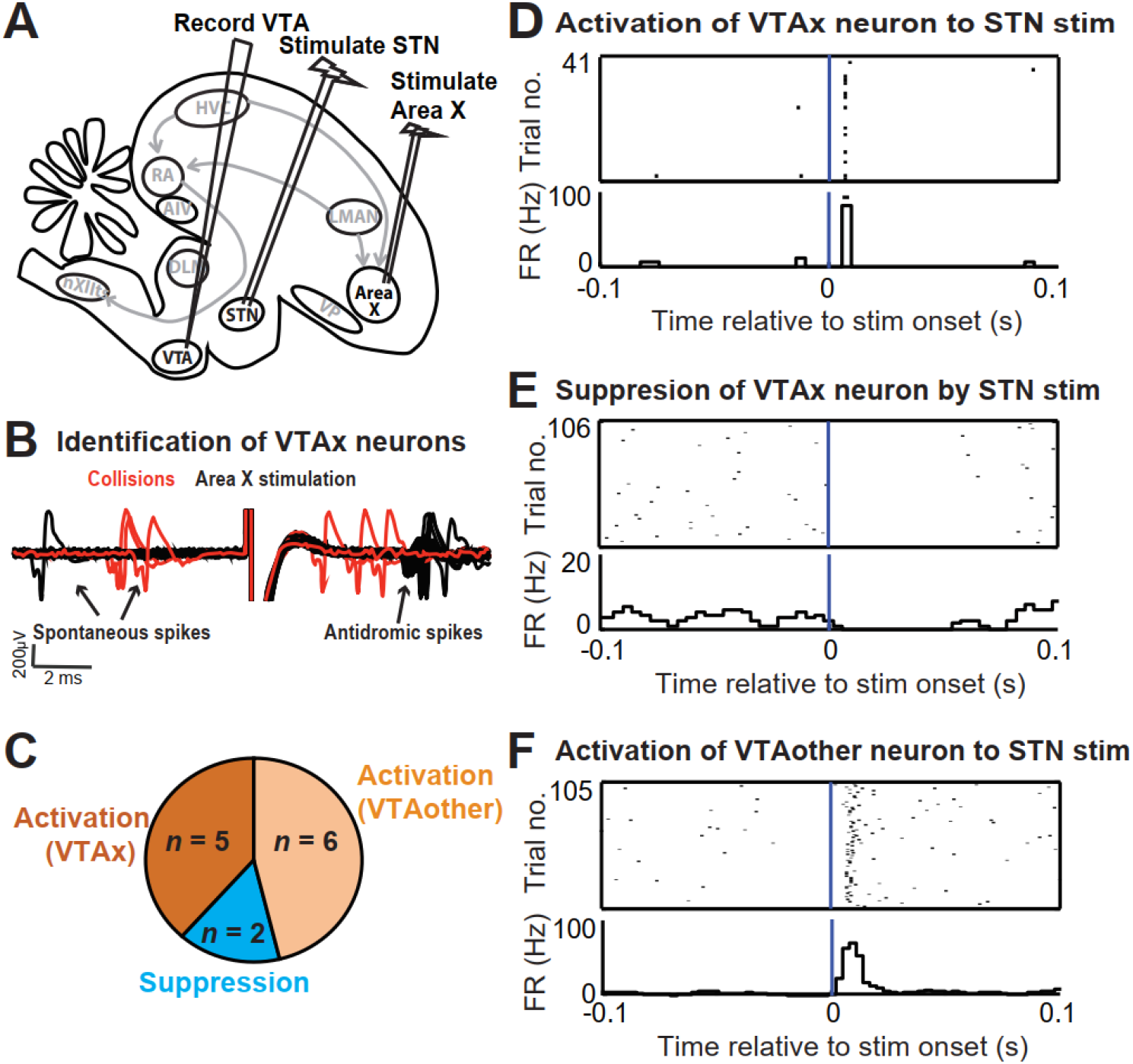
STN stimulation activates VTA neurons including Area X-projecting VTA neurons. A. Bipolar electrodes were implanted in AreaX and STN in anaesthetized birds to test the effect of STN microstimulation on the firing of VTA neurons. B. All VTA neurons recorded were tested to see if they were AreaX-projecting neurons (VTAx) by collision testing of spontaneous spikes with antidromic spikes generated by AreaX stimulation. VTA neurons were then classified as either X-projectors (VTAx) or non-projectors (VTAother). C. A total of 13 VTA neurons were tested, out of which 7 were identified as VTAx. 5 VTAx neurons were activated by STN stimulation, while 2 were suppressed. All VTAother neurons showed activation D. Example of a VTAx neuron exhibiting a short latency (4 ms) activation to STN stimulation. E. Example of a VTAx neuron suppressed by STN stim. F. Example of a VTAother neuron activated by STN stimulation at a latency of 5.5 ms

### A small fraction of STN neurons exhibit song-timing and performance error signals in singing birds

We next wondered if STN neurons exhibit any singing- and/or error-related signals. To test this, we recorded 62 neurons in 7 (out of 36 implanted) birds while we controlled perceived error with syllable-targeted DAF, as previously described (Chen et al. 2019; Gadagkar et al. 2016). In 4/7 birds, we also recorded movements with head-mounted accelerometers. To test for singingrelated changes in firing, we examined for significant firing rate changes during singing, for syllable-locked timing, and for DAF-associated error responses. Most STN neurons were not singing related by these criteria (51/62). However, a small population of STN neurons (*n*=11 out of 62 recorded) in singing birds exhibited time-locked firing during singing (*n*=9) and performance error signaling (*n*=2). The firing rate of all the time-locked neurons during singing exhibited high variability ranging from 2.3 Hz to 92 Hz. Out of the 9 song-locked neurons, 4 neurons exhibited a high peak firing rate during singing (> 100 Hz) and mean firing rate (during singing) 59.7 ± 23.3 Hz (mean ± sd). They also exhibited a large increase in firing at the onset of motif and then precise time-locked firing within the song (example in Fig. 3A). 4 other neurons exhibited mostly low firing rates to almost no firing outside of singing and precise time-locked firing with increase in firing rate during singing (mean ± sd, 7.3 ± 4.7 Hz) (example in Fig. 3B). One of the neurons exhibited a precise, sparse firing during singing (Fig. 3C). None of these neurons exhibited any difference in firing rates between distorted and undistorted trials. All these song timing neurons exhibited a significant difference in firing rate during singing compared to silent periods (*n*=9, *p* < 0.05, Wilcoxon signed rank test) (blue circles, Fig. 1F).

**Fig. 3.**
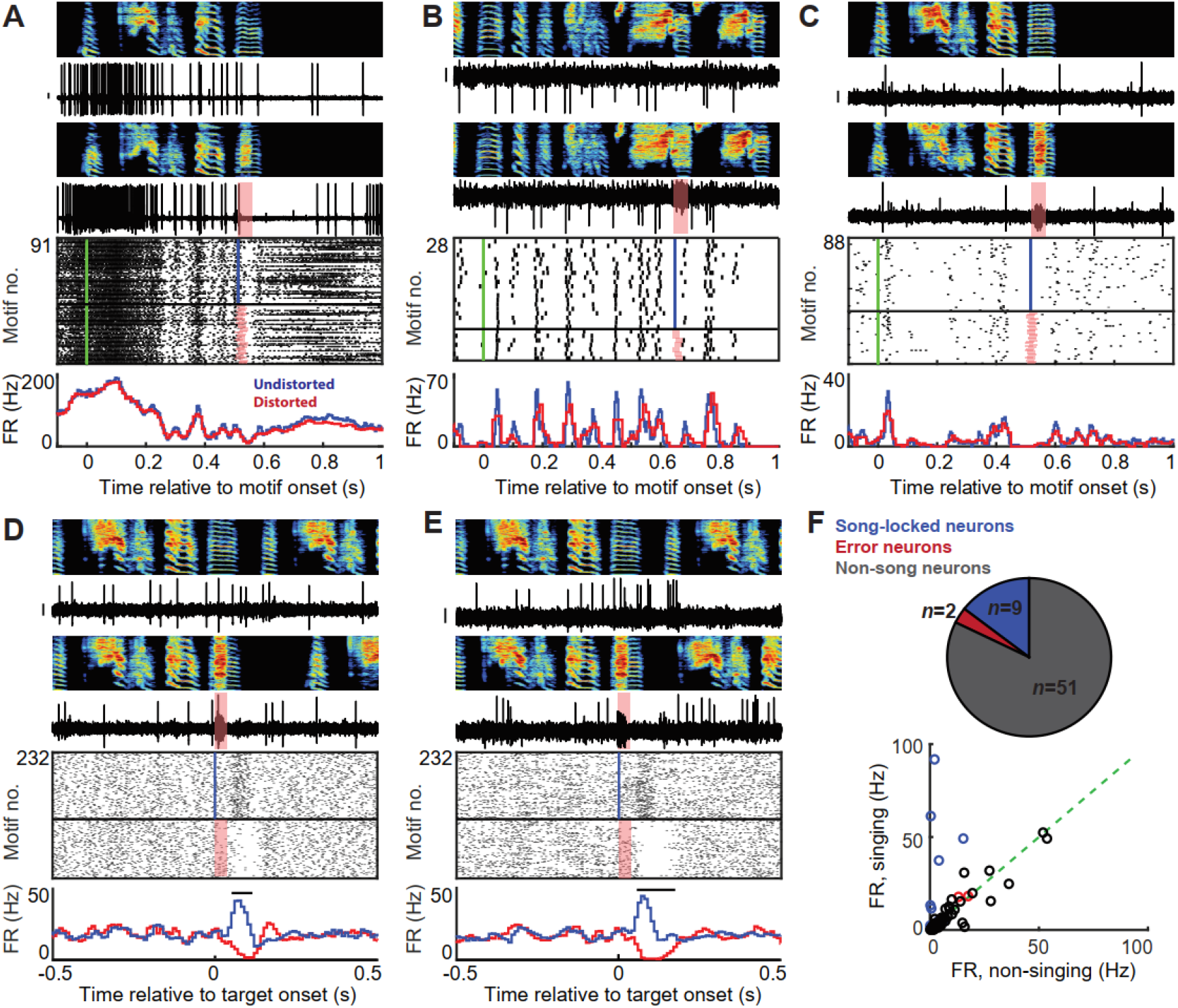
A small population of STN neurons exhibit song-timing and performance error signals. A—E. Examples of STN neurons in singing birds are shown with distorted and undistorted motifs on top along with the corresponding neural traces. For each neuron, rasters and firing rate histograms are depicted with distorted renditions in red and undistorted renditions in blue. Scale bar on all neural traces is 0.1 mV. A—C. Example neurons exhibiting varying firing rates and precise time-locked firing during song. The motif aligned rasters and firing rate histograms indicate no difference in neural response between distorted and undistorted trials. The green line in the rasters indicates motif onset time. D—E. Two of the STN neurons recorded exhibited performance error signal, with decrease in firing rate on distorted trials and increase in firing rate during undistorted trials, as indicated by the target-aligned rasters and firing rate histograms. F. Overall, a small fraction of all neurons recorded exhibited song-locked firing rate modulation or error response. The song timing neurons exhibited increased firing rate during song compared to outside song (n=8/9) (top right panel). The error neurons and the other neurons did not exhibit a significant difference in firing rate during versus outside song.

The precision of time-locked firing was measured by computing the inter-motif correlation coefficient (IMCC) (Chen et al. 2019; Kao et al. 2008; Ölveczky et al. 2005). Significance of the IMCC values were assessed by generating ‘new’ IMCC values from shuffled data (random, circular time shifted spike trains) and a Kolmogorov-Smirnov test was used to compare the actual IMCC value to the ‘new’ IMCC value of the shuffled data (Chen et al. 2019). All 9 neurons exhibited significant IMCC values (mean ± sd, 0.25 ± 0.26; range, 0.04 to 0.76, *p* < 0.001). The high firing rate neurons exhibited a higher value of IMCC. Two STN neurons exhibited performance error signals with suppression of firing following DAF during distorted renditions and increased firing following absence of DAF during undistorted renditions (Fig. 3D,E), similar to VTAx DA neurons (Gadagkar et al. 2016). Like VTAx neurons, they did not exhibit any significant song-locked activity (IMCC < 0.01, *p* > 0.05) and did not exhibit a significant difference in firing rates during and outside song (*p* > 0.05, Wilcoxon signed rank test) (red circles, Fig. 3F). Singing and non-singing-related neurons were spatially intermingled (Fig. 5A).

### Songbird STN neurons exhibit movement-related discharge

A total of 36 neurons were recorded from birds (*n*=4) implanted with headstages mounted with accelerometers, facilitating the assessment of movement-related activity in the finch STN. Out of these neurons, 20 showed significant firing rate changes following movement onsets/offsets and 8 of these showed movement onset aligned firing rate increase only outside singing. The firing rates of these movement neurons ranged from 1 Hz to 50 Hz with a mean firing rate outside singing 9.12 ± 12.6 Hz (mean ± sd). 14 neurons that exhibited movement onset firing rate changes during both singing and non-singing periods (example, Fig. 4A) did not have any significant difference between their firing rates outside singing versus during singing (*p* > 0.05, Wilcoxon signed rank test) (Fig.4F). The other 8 neurons had significantly different (*p* < 0.05, Wilcoxon signed rank test) firing rates outside singing (mean ± sd, 4.37 ± 4.8 Hz) compared to during singing (mean ± sd, 1.51 ± 0.9 Hz) and exhibited movement onset firing rate increase only outside singing (example, Fig. 4B). A heat map of the z-scored firing rate change for all neurons aligned to movement onset showed a variability in latency to response, but most of the response was following movement onset (Fig. 4C, D). The latency to movement onset varied from −15 ms to 55 ms (Fig. 4G) with a median latency of 15 ms. 4 of these neurons exhibited a movement onset aligned suppression of firing rate. Our results showing movement aligned firing rate modulation of finch STN neurons is consistent with the known motor functions of the mammalian STN (Fife et al. 2017; Georgopoulos et al. 1983; Hamani et al. 2004). In addition, neurons in other areas of the finch brain, namely VTA and VP have been shown to contain neurons that encode movement and display differential encoding of movement depending on the behavioural state of the animal (singing versus nonsinging) (Chen et al. 2021). Our results suggest that the finch STN also contains a population of neurons encoding movement independent of state and another smaller population that exhibits singing-related gating of movement signaling. None of the movement related neurons exhibited any song-locked firing rate modulation according to the criteria defined previously (IMCC < 0.01, *p* > 0.05).

**Fig. 4.**
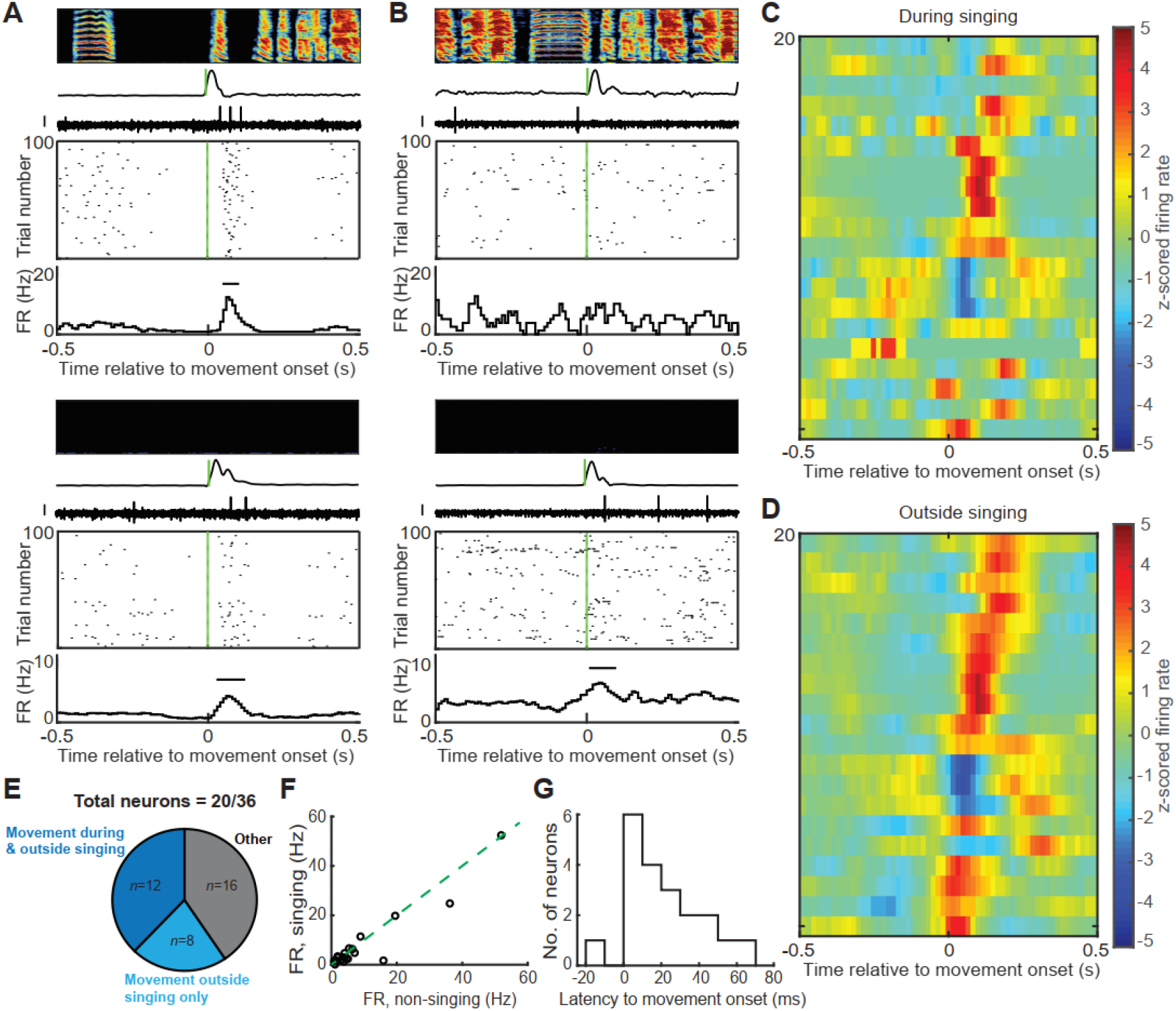
Songbird STN neurons exhibit movement aligned firing rate changes. A. Example of a neuron that exhibited movement onset increase in firing rate both during and outside singing. Top panel shows the spectrogram of the song with time-aligned smoothed rectified accelerometer signal and neural trace. The corresponding raster plot is shown below with the firing rate (FR in Hz) histogram. The bar in the histogram depicts significant change in firing rate (z-test, p < 0.05). Bottom panel shows the raster plot and firing rate histogram along with the neural trace and accelerometer signal corresponding to non-singing period. Scale bar on all neural traces is 0.1 mV. B. Example of a neuron that exhibited no movement aligned firing rate changes during song (top panel) and movement onset firing rate increase during non-singing period (bottom panel). C, D. The z-scored firing rate change was plotted for all neurons (n=20) during singing (C) and outside singing (D). 3 neurons were suppressed following movement onset, while the others showed increased firing following movement onset with a response duration in the range of hundreds of ms. E. A total of 36 neurons were recorded from birds with headstages mounted with accelerometers. Out of these, 20 neurons exhibited significant movement aligned firing rate changes and 12 of these did so during both singing and non-singing periods. 8 neurons were movement aligned only outside singing. F. Most of the neurons exhibited low firing rates and did not show any significant difference between firing rate during singing and outside singing. G. Latencies to movement onset were computed based on the window of significant firing rate change in the firing rate histogram. A histogram of the latencies has been plotted. Latency values ranged from −15 ms to 65 ms, with the majority of the neurons exhibiting significant firing rate change following movement onset.

## Discussion

The STN is an evolutionarily conserved, glutamatergic nucleus in the vertebrate diencephalon integrated into the indirect and hyperdirect pathways of the BG, where it is thought to serve a primary role of halting movements (Fife et al. 2017; Frank 2006; Nambu et al. 2002; Schmidt et al. 2013; Wessel et al. 2016). The STN features most prominently as the subject of pathophysiological studies and has, over the years, emerged as the singular therapeutic focus in patients with Parkinson’s disease (Benarroch 2008; Frank et al. 2007; Hamani et al. 2004). More recently the STN has been postulated to serve a role beyond that of a simple ‘motor brake’ (Breysse et al. 2015; Lardeux et al. 2009; Temel et al. 2005). First, STN neurons project directly to midbrain DA neurons, and provide their main source of glutamatergic input (Ogawa et al. 2014; Watabe-Uchida et al. 2012). Second, STN stimulation directly activates DA neurons and increases striatal DA levels (Bruet et al. 2001; Meissner et al. 2002, 2003; Paul et al. 2000). Third, STN neurons exhibit reward- and reward cue-related signals (Baunez et al. 2002; Breysse et al. 2015; Tian et al. 2016). Finally, STN lesions in mammals impair reward-seeking behavior in animals and deep brain stimulation of STN in Parkinsonian patients have demonstrated to ameliorate both motor and cognitive deficits (Baunez et al. 2002, 2005; Van Wouwe et al. 2011). These studies in mammals that illustrate functional connections between STN and dopaminergic midbrain led us to hypothesize that STN could be playing a hitherto undiscovered role in song learning, which is known to rely on DA evaluation signals.

By combining anatomical and functional circuit mapping and electrophysiology in singing birds, we discovered that the STN is interconnected with AIV, VP, and VTA, three structures known to play a role in song evaluation. We also discovered that a small fraction of STN exhibits singing related signals, including error signals. This first-ever investigation into the functions of songbird STN suggests a possible role in song evaluation and expands our understanding of the singing-related portions of the songbird brain beyond the classic song system. These findings also place the songbird STN into a similar circuit architecture known in mammals. While our findings support the idea that STN is part of the song evaluation system upstream VTA, lesion studies that would firmly test the causal role of STN in song learning were unfortunately not possible. We attempted to bilaterally lesion STN to assess its role in song learning but 4/4 lesioned birds exhibited such dramatic movement impairments post-operatively that they were euthanized. Such severe post-lesion motor impairments prevent the assessment of any other learning (song learning in this case) deficits. Future studies using cell type-specific viral targeting methods to specifically ablate or optogenetically regulate VTA-projecting STN neurons may be able to causally test the necessity of this projection for DA error signaling and song learning (Fig. 5B). A second caveat is that only a small fraction of STN neurons exhibited singing- or error-related activity (11/62). Most STN neurons exhibited movement-locked activity, yet singing- and movement-related neurons were spatially intermingled. Thus, unlike the nuclei of the song system, where all neurons are singing-related, the songbird STN may serve other more general motor evaluation functions. Notably, a similar intermixing of singing- and non-singing related signals were observed in VP and VTA, and both of these structures exhibited a similarly small fraction of neurons that encoded song timing or error (Chen et al. 2019, 2021; Gadagkar et al. 2016). A key feature of the DA evaluation signals in singing birds was that the activations occurred when a predicted error did not occur. In contrast, during reward seeking, activations are driven by explicit sensory events such as rewards or cues that predict them. Our study raises the possibility that precisely timed DA activations associated with better-than-predicted performances in mammals could depend on the STN-DA projection, which could be tested in future studies where motor performances are carried out.

**Fig. 5.**
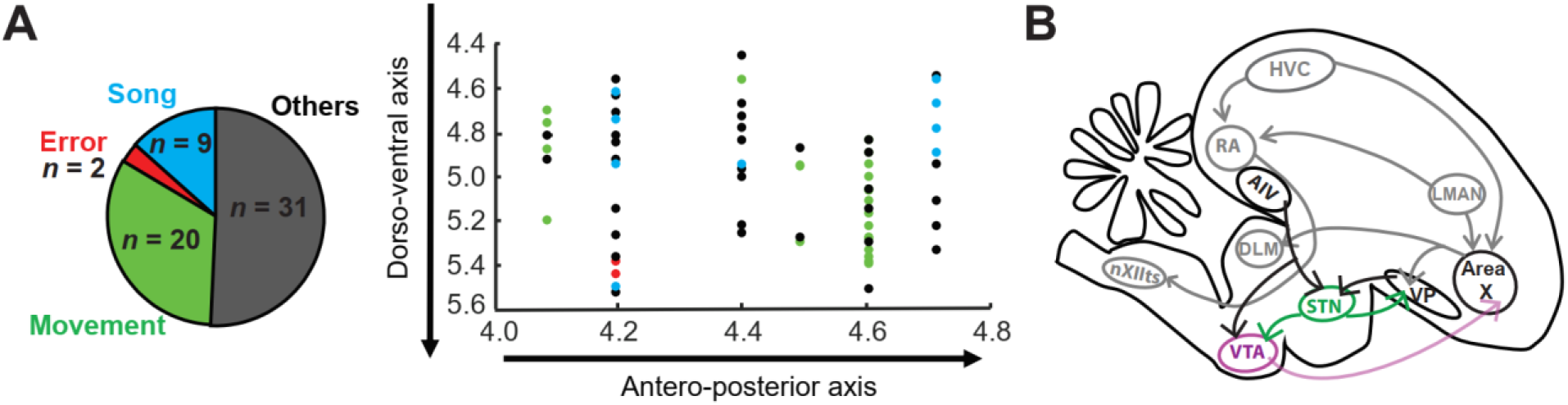
STN in zebra finches. A. We recorded a total of 62 STN neurons in singing birds and found that they were spatially distributed with no functional clustering. B. In bold are highlighted anatomical connections that we confirmed in our experiments (AIV to STN with collateral to VTA, VP to STN, STN to VP and STN to VTA). We suggest future experiments to specifically target VTAx-projecting STN neurons to assess how suppressing or ablating this population of neurons would alter the error signalling in VTAx neurons.

## Acknowledgement

This work was supported by funding from NIH R01NS094667. We thank Archana Podury for help with anatomy and members of the Goldberg lab for comments on the manuscript.

## Notes

### Competing Interest Statement

The authors have declared no competing interest.

## References

Adamantidis AR, Tsai H-C, Boutrel B, Zhang F, Stuber GD, Budygin EA, Touriño C, Bonci A, Deisseroth K, Lecea L de. Optogenetic Interrogation of Dopaminergic Modulation of the Multiple Phases of Reward-Seeking Behavior. J Neurosci 31: 10829–10835, 2011.

Andalman AS, Fee MS. A basal ganglia-forebrain circuit in the songbird biases motor output to avoid vocal errors. PNAS 106: 12518–12523, 2009.

Baunez C, Amalric M, Robbins TW. Enhanced food-related motivation after bilateral lesions of the subthalamic nucleus. J Neurosci 22: 562–568, 2002.

Baunez C, Dias C, Cador M, Amalric M. The subthalamic nucleus exerts opposite control on cocaine and “natural” rewards. Nat Neurosci 8: 484–489, 2005.

Benarroch EE. Subthalamic nucleus and its connections: Anatomic substrate for the network effects of deep brain stimulation. Neurology 70: 1991–1995, 2008.

Breysse E, Pelloux Y, Baunez C. The Good and Bad Differentially Encoded within the Subthalamic Nucleus in Rats. eNeuro 2, 2015.

Bromberg-Martin ES, Matsumoto M, Hikosaka O. Dopamine in Motivational Control: Rewarding, Aversive, and Alerting. Neuron 68: 815–834, 2010.

Bruet N, Windels F, Bertrand A, Feuerstein C, Poupard A, Savasta M. High Frequency Stimulation of the Subthalamic Nucleus Increases the Extracellular Contents of Striatal Dopamine in Normal and Partially Dopaminergic Denervated Rats. Journal of Neuropathology & Experimental Neurology 60: 15–24, 2001.

Carouso-Peck S, Goldstein MH. Female Social Feedback Reveals Non-imitative Mechanisms of Vocal Learning in Zebra Finches. Current Biology 29: 631–636.e3, 2019.

Chen R, Gadagkar V, Roeser AC, Puzerey PA, Goldberg JH. Movement signaling in ventral pallidum and dopaminergic midbrain is gated by behavioral state in singing birds. bioRxiv 2020.06.22.164814, 2021.

Chen R, Goldberg JH. Actor-critic reinforcement learning in the songbird. Current Opinion in Neurobiology 65: 1–9, 2020.

Chen R, Puzerey PA, Roeser AC, Riccelli TE, Podury A, Maher K, Farhang AR, Goldberg JH. Songbird Ventral Pallidum Sends Diverse Performance Error Signals to Dopaminergic Midbrain. Neuron 103: 266–276, 2019.

Colquitt BM, Merullo DP, Konopka G, Roberts TF, Brainard MS. Cellular transcriptomics reveals evolutionary identities of songbird vocal circuits. Science 371, 2021.

Darbaky Y, Baunez C, Arecchi P, Legallet E, Apicella P. Reward-related neuronal activity in the subthalamic nucleus of the monkey. Neuroreport 16: 1241–1244, 2005.

Doupe AJ, Kuhl PK. BIRDSONG AND HUMAN SPEECH: Common Themes and Mechanisms. Annual Review of Neuroscience 22: 567–631, 1999.

Doupe AJ, Perkel DJ, Reiner A, Stern EA. Birdbrains could teach basal ganglia research a new song. Trends Neurosci 28: 353–363, 2005.

Doya K, Sejnowski TJ. A Computational Model of Birdsong Learning by Auditory Experience and Auditory Feedback. In: Central Auditory Processing and Neural Modeling, edited by Poon PWF, Brugge JF. Springer US, p. 77–88.

Fee MS, Goldberg JH. A hypothesis for basal ganglia-dependent reinforcement learning in the songbird. Neuroscience 198: 152–170, 2011.

Fife KH, Gutierrez-Reed NA, Zell V, Bailly J, Lewis CM, Aron AR, Hnasko TS. Causal role for the subthalamic nucleus in interrupting behavior. Elife 6, 2017.

Frank MJ. Hold your horses: a dynamic computational role for the subthalamic nucleus in decision making. Neural Netw 19: 1120–1136, 2006.

Frank MJ, Samanta J, Moustafa AA, Sherman SJ. Hold your horses: impulsivity, deep brain stimulation, and medication in parkinsonism. Science 318: 1309–1312, 2007.

Gadagkar V, Puzerey PA, Chen R, Baird-Daniel E, Farhang AR, Goldberg JH. Dopamine neurons encode performance error in singing birds. Science 354: 1278–1282, 2016.

Gadagkar V, Puzerey PA, Goldberg JH. Dopamine neurons change their tuning according to courtship context in singing birds. bioRxiv 822817, 2019.

Gale SD, Person AL, Perkel DJ. A novel basal ganglia pathway forms a loop linking a vocal learning circuit with its dopaminergic input. Journal of Comparative Neurology 508: 824–839, 2008.

Georgopoulos AP, DeLong MR, Crutcher MD. Relations between parameters of step-tracking movements and single cell discharge in the globus pallidus and subthalamic nucleus of the behaving monkey. J Neurosci 3: 1586–1598, 1983.

Grillner S, Robertson B, Stephenson-Jones M. The evolutionary origin of the vertebrate basal ganglia and its role in action selection. The Journal of Physiology 591: 5425–5431, 2013.

Groenewegen HJ, Berendse HW, Haber SN. Organization of the output of the ventral striatopallidal system in the rat: ventral pallidal efferents. Neuroscience 57: 113–142, 1993.

Haber SN, Lynd-Balta E, Mitchell SJ. The organization of the descending ventral pallidal projections in the monkey. J Comp Neurol 329: 111–128, 1993.

Hahnloser R, Ganguli S. Vocal learning with inverse models. In: Principles of Neural Coding, edited by Panzeri S, Quiroga P. 2013, p. 547–564.

Hamani C, Saint-Cyr JA, Fraser J, Kaplitt M, Lozano AM. The subthalamic nucleus in the context of movement disorders. Brain 127: 4–20, 2004.

Hisey E, Kearney MG, Mooney R. A common neural circuit mechanism for internally guided and externally reinforced forms of motor learning. Nat Neurosci 21: 589–597, 2018.

Hoffmann LA, Saravanan V, Wood AN, He L, Sober SJ. Dopaminergic Contributions to Vocal Learning. J Neurosci 36: 2176–2189, 2016.

Jiao Y, Medina L, Veenman CL, Toledo C, Puelles L, Reiner A. Identification of the anterior nucleus of the ansa lenticularis in birds as the homolog of the mammalian subthalamic nucleus. J Neurosci 20: 6998–7010, 2000.

Joel D, Weiner I. The connections of the primate subthalamic nucleus: indirect pathways and the open-interconnected scheme of basal ganglia-thalamocortical circuitry. Brain Res Brain Res Rev 23: 62–78, 1997.

Kao MH, Wright BD, Doupe AJ. Neurons in a Forebrain Nucleus Required for Vocal Plasticity Rapidly Switch between Precise Firing and Variable Bursting Depending on Social Context. J Neurosci 28: 13232–13247, 2008.

Kearney MG, Warren TL, Hisey E, Qi J, Mooney R. Discrete Evaluative and Premotor Circuits Enable Vocal Learning in Songbirds. Neuron 104: 559–575.e6, 2019.

Kooy DVD, Hattori T. Single subthalamic nucleus neurons project to both the globus pallidus and substantia nigra in rat. Journal of Comparative Neurology 192: 751–768, 1980.

Lardeux S, Pernaud R, Paleressompoulle D, Baunez C. Beyond the reward pathway: coding reward magnitude and error in the rat subthalamic nucleus. J Neurophysiol 102: 2526–2537, 2009.

Mandelblat-Cerf Y, Las L, Denisenko N, Fee MS. A role for descending auditory cortical projections in songbird vocal learning. eLife 3: e02152, 2014.

Marler P. Three models of song learning: Evidence from behavior. Journal of Neurobiology 33: 501–516, 1997.

Meissner W, Harnack D, Paul G, Reum T, Sohr R, Morgenstern R, Kupsch A. Deep brain stimulation of subthalamic neurons increases striatal dopamine metabolism and induces contralateral circling in freely moving 6-hydroxydopamine-lesioned rats. Neuroscience Letters 328: 105–108, 2002.

Meissner W, Harnack D, Reese R, Paul G, Reum T, Ansorge M, Kusserow H, Winter C, Morgenstern R, Kupsch A. High-frequency stimulation of the subthalamic nucleus enhances striatal dopamine release and metabolism in rats. Journal of Neurochemistry 85: 601–609, 2003.

Mello CV, Vates E, Okuhata S, Nottebohm F. Descending auditory pathways in the adult male zebra finch (Taeniopygia Guttata). Journal of Comparative Neurology 395: 137–160, 1998.

Murphy K, James LS, Sakata JT, Prather JF. Advantages of comparative studies in songbirds to understand the neural basis of sensorimotor integration. Journal of Neurophysiology 118: 800–816, 2017.

Nambu A, Tokuno H, Takada M. Functional significance of the cortico-subthalamo-pallidal “hyperdirect” pathway. Neurosci Res 43: 111–117, 2002.

Ogawa SK, Cohen JY, Hwang D, Uchida N, Watabe-Uchida M. Organization of Monosynaptic Inputs to the Serotonin and Dopamine Neuromodulatory Systems. Cell Reports 8: 1105–1118, 2014.

Ölveczky BP, Andalman AS, Fee MS. Vocal Experimentation in the Juvenile Songbird Requires a Basal Ganglia Circuit. PLOS Biology 3: e153, 2005.

Paul G, Reum T, Meissner W, Marburger A, Sohr R, Morgenstern R, Kupsch A. High frequency stimulation of the subthalamic nucleus infuences striatal dopaminergic metabolism in the naive rat. NeuroReport 11: 441–444, 2000.

Paul K, Johnson SW. Post-inhibitory rebound properties of dopaminergic cells of the ventral tegmental area. Neuroscience Research Communications 33: 147–157, 2003.

Person AL, Gale SD, Farries MA, Perkel DJ. Organization of the songbird basal ganglia, including area X. J Comp Neurol 508: 840–866, 2008.

Reiner A, Medina L, Veenman CL. Structural and functional evolution of the basal ganglia in vertebrates. Brain Research Reviews 28: 235–285, 1998.

Schmidt R, Leventhal DK, Mallet N, Chen F, Berke JD. Canceling actions involves a race between basal ganglia pathways. Nat Neurosci 16: 1118–1124, 2013.

Schultz W. Predictive reward signal of dopamine neurons. J Neurophysiol 80: 1–27, 1998.

Schultz W, Dayan P, Montague PR. A Neural Substrate of Prediction and Reward. Science 275: 1593–1599, 1997.

Sohrabji F, Nordeen EJ, Nordeen KW. Selective impairment of song learning following lesions of a forebrain nucleus in the juvenile zebra finch. Behavioral and Neural Biology 53: 51–63, 1990.

Sutton RS, Barto AG. Reinforcement Learning, second edition: An Introduction. MIT Press, 2018.

Tchernichovski O, Mitra PP, Lints T, Nottebohm F. Dynamics of the Vocal Imitation Process: How a Zebra Finch Learns Its Song. Science 291: 2564–2569, 2001.

Temel Y, Blokland A, Steinbusch HW, Visser-Vandewalle V. The functional role of the subthalamic nucleus in cognitive and limbic circuits. Prog Neurobiol 76: 393–413, 2005.

Tian J, Huang R, Cohen JY, Osakada F, Kobak D, Machens CK, Callaway EM, Uchida N, Watabe-Uchida M. Distributed and Mixed Information in Monosynaptic Inputs to Dopamine Neurons. Neuron 91: 1374–1389, 2016.

Tsai HC, Zhang F, Adamantidis A, Stuber GD, Bonci A, de Lecea L, Deisseroth K. Phasic firing in dopaminergic neurons is sufficient for behavioral conditioning. Science 324: 1080–1084, 2009.

Tumer EC, Brainard MS. Performance variability enables adaptive plasticity of ‘crystallized’ adult birdsong. Nature 450: 1240–1244, 2007.

Van Wouwe NC, Wylie SA, Van Den Wildenberg WPM, Band GPH, Abisogun AA, Elias WJ, Frysinger RC, Ridderinkhof RKR. Deep Brain Stimulation of the Subthalamic Nucleus Improves Reward-Based Decision-Learning in Parkinson’s Disease. Front Hum Neurosci 5, 2011.

Watabe-Uchida M, Zhu L, Ogawa SK, Vamanrao A, Uchida N. Whole-Brain Mapping of Direct Inputs to Midbrain Dopamine Neurons. Neuron 74: 858–873, 2012.

Wessel JR, Jenkinson N, Brittain J-S, Voets SHEM, Aziz TZ, Aron AR. Surprise disrupts cognition via a fronto-basal ganglia suppressive mechanism. Nat Commun 7: 11195, 2016.

Woolley SC. Dopaminergic regulation of vocal-motor plasticity and performance. Current Opinion in Neurobiology 54: 127–133, 2019.

Xiao L, Chattree G, Oscos FG, Cao M, Wanat MJ, Roberts TF. A Basal Ganglia Circuit Sufficient to Guide Birdsong Learning. Neuron 98: 208–221.e5, 2018.

Zweifel LS, Parker JG, Lobb CJ, Rainwater A, Wall VZ, Fadok JP, Darvas M, Kim MJ, Mizumori SJ, Paladini CA, Phillips PE, Palmiter RD. Disruption of NMDAR-dependent burst firing by dopamine neurons provides selective assessment of phasic dopamine-dependent behavior. Proc Natl Acad Sci U S A 106: 7281–7288, 2009.

